# Synthesis and evaluation of novel *N,N*-dialkylcinnamic acid-based mitochondrial pyruvate carrier inhibitors: Biosynthetic and energetic lethality of targeting metabolic plasticity in cancer

**DOI:** 10.64898/2026.01.12.699093

**Authors:** Sravan Jonnalagadda, Conor T. Ronayne, Shirisha Jonnalagadda, Zachary. S. Gardner, Venkatram R. Mereddy

**Affiliations:** Integrated Biosciences Graduate Program, University of Minnesota, Duluth, MN 55812; Department of Pharmacy Practice & Pharmaceutical Sciences, University of Minnesota, Duluth, MN 55812; Department of Chemistry and Biochemistry, University of Minnesota Duluth, Duluth, MN 55812; Department of Chemistry, University of Wisconsin-Whitewater, Whitewater, WI 53190

**Keywords:** mitochondrial pyruvate carrier, monocarboxylate transporter, metabolic plasticity, tumor metabolism, anticancer drugs

## Abstract

Novel functionalized cyanocinnamic acid based MPC inhibitors based on pharmacologically privileged *N*-piperazinyl and *N-*piperidinyl drug templates have been synthesized for potential cancer treatment. *In vitro* cell proliferation inhibition studies with these derivatives **2-4** show activity in the low micromolar range. Seahorse XFe96 based mitochondrial stress tests also illustrate the ability of **2-4** to potently and acutely inhibit numerous parameters of mitochondrial respiration in MDA-MB-231, WiDr, and 4T1 cells. Further analyses of the lead compound **3** in permeabilized 4T1 cells provide evidence of specific inhibition of pyruvate driven respiration without affecting glutamate or succinate fueled respiratory processes. Combination studies with GLUT1 inhibitor BAY-876 illustrate the capacity of compound **3** to inhibit metabolic plasticity in triple negative breast cancer MDA-MB-231 cells and is synergistic in inhibiting cell proliferation in aggressive stage IV breast cancer 4T1.

## INTRODUCTION

The metabolic phenotype of cancer cells is generally different from that of normal cells due to elevated demand for energy and anabolic precursors to sustain rapid proliferation.^1,2^ Metabolic reprogramming is one of the critical hallmarks of cancer progression. Tumor cells frequently exhibit a heightened glycolytic flux, even under normoxic conditions and persistent glycolytic activity is not only critical for ATP generation but also underpins resistance to apoptosis, enhanced invasiveness, and metastatic potential.^3^ Glycolytic intermediates also feed into ancillary biosynthetic routes, including the pentose phosphate pathway for de novo nucleotide synthesis. Consequently, glycolytic inhibition precipitates energetic insufficiency and depletion of biosynthetic precursors.

Recent investigations have further illuminated the complexity of cancer metabolism, emphasizing the contribution of mitochondrial oxidative phosphorylation (OxPhos) to tumor bioenergetics and survival.^2,4,5^ Many tumor cells retain substantial mitochondrial function, with OxPhos generating a significant fraction of cellular ATP and tricarboxylic acid (TCA) cycle intermediates provide essential substrates for lipid, amino acid, and nucleotide biosynthesis. Inhibition of OxPhos tin cancer cells therefore results in profound ATP depletion and TCA cycle dysfunction, depriving cancer cells of critical anabolic substrates required for growth and survival. Tumor microenvironments characterized by fluctuating oxygen and nutrient availability drive dynamic switching between glycolytic and oxidative phenotypes—a metabolic plasticity that enables tumor persistence under hostile conditions.^6-7^ Accordingly, dual inhibition of glycolysis and OxPhos represents a promising therapeutic strategy to simultaneously disrupt energy production and biosynthetic capacity, thereby suppressing tumor progression.

The mitochondrial pyruvate carrier (MPC) mediates the import of cytosolic pyruvate into the mitochondrial matrix.^8-9^ Situated within the inner mitochondrial membrane, the MPC integrates glycolytic and mitochondrial metabolism by channeling pyruvate into the TCA cycle.^8-9^ Through this coupling of glycolysis and OxPhos, the MPC fulfills both energetic and anabolic requirements of proliferating tumor cells. Recent studies reveal that oxidative tumor phenotypes exhibit enhanced mitochondrial respiration and biosynthetic activity, and pharmacologic inhibition of MPC has demonstrated efficacy in multiple preclinical tumor models.^10^ The MPC is a relatively understudied in clinical oncology and occupies a central regulatory position in coordinating cytosolic and mitochondrial metabolism and in modulating monocarboxylate transporter (MCT)-dependent metabolite flux. As such, it represents a compelling target for therapeutic intervention.

Monocarboxylate transporters (MCTs) belong to the solute carrier 16 (SLC16) family, which comprises 14 known isoforms.^14^ Among these, MCT1–4 facilitate proton-linked translocation of monocarboxylates, including lactate, pyruvate, and select ketone bodies.^15-20^ Our lab has developed a wide variety of potent MCT1/4 based on *N,N*-dialkylcyanocinnamic acid templates^18,20^. These drug candidates have been evaluated on a variety of cancer cell lines and xenograft models where they exhibited significant tumor growth reduction as single agents and in combination with current chemotherapeutics. The known MPC inhibitor UK5099 contains a cyanocinnamic acid unit, and we sought to explore the potential of our lead *N,N*-dialkylcyanocinnamic acid candidates^18,20^ as MPC inhibitors. Here, we find that *N,N*-dialkylcyanocinnamic acid compounds inhibit the MPC potently. Further, chemical modification of the *N,N*-dialkylcyanocinnamic acid template onto diverse and robust pharmaceutical templates has provided new generation MPC inhibitors with enhanced cancer cell proliferation inhibition properties while retaining MPC inhibition properties. Finally, we explore the ability of lead MPC inhibitor **3** -to synergize with GLUT1 inhibitors to arrest compensatory metabolic plasticity and cancer cell proliferation.

## RESULTS

### N,N-dialkylcyanocinnamic acid derivatives inhibit the mitochondrial pyruvate carrier

We have previously reported the synthesis and evaluation of *N,N*-dialkylcyanocinnamic acid derivatives **1a** and **1b** with potent MCT1/4 inhibition properties (**Figure 1A**)^18, 20^. Literature reports suggest that inhibition of mitochondrial pyruvate flux with *N,N-*dialkyl aminocarboxycoumarin derivatives (ex. 7ACC2, **Figure 1A**) leads to increased cytosolic pyruvate concentrations that can modulate MCT-mediated lactate uptake, where these derivatives were defined as MPC inhibitors^21^. In fact, we also reported^19^ on *N,N-*dialkyl aminocarboxycoumarin derivatives that exhibited MCT1 inhibition properties, consistent with feedback-mediated inhibition as reported above^21^. In line with these findings, we have recently reported on the synthesis and evaluation of novel *N,N*-dialkyl aminocarboxycoumarin derivatives that potently inhibit MPC function and arrest tumor growth in mouse models of MCT1 expressing breast cancers^22^. Based on pharmacological similarities between coumarin and cyanocinnamic acid derivatives (**Figure 1A**), we sought to explore the ability of our first-generation *N,N*-dialkylcyanocinnamic MCT inhibitors to also inhibit the MPC. Here, using our previously reported Seahorse XFe-based MPC inhibition assay^22^, we evaluated the ability of **1a** and **1b** to inhibit pyruvate driven respiration in permeabilized 4T1 breast cancer cells. Our results illustrated that **1a** and **1b** inhibit MPC function to a similar magnitude of known MPC inhibitors UK5099 and 7ACC2 when tested at similar concentrations (1 µM, **Figure 1B-C**). Dose-response curves indicate that **1a** and **1b** inhibit pyruvate driven respiration at 133 and 38 nM, respectively (**Figure 1D-E**). Although **1a** and **1b** are potent MPC inhibition properties, these compounds are limited with pharmaceutical shortcomings. We previously evaluated the metabolic stability of compound **1a** and this derivative showed limited metabolic stability with low biological half-lives and rapid clearance rates^18^. To further improve its metabolic stability, we envisioned to modify the first generation *N,N*-dialkylcyanocinnamic acid template with pharmacologically privileged templates with further structural diversity and drug-like properties based on cetirizine [(*S*)-1-((4-chlorophenyl)(phenyl)methyl)piperazine], quetiapine [11-(piperazin-1-yl)dibenzo[b,f][1,4]thiazepine] and paroxetine [(3*S*,4*R*)-3-((benzo[d][1,3]dioxol-5-yloxy)methyl)-4-(4-fluorophenyl)piperidine] to synthesize cyanocinnamic acids (**2-4, Figure 1F**).

**Figure 1.**
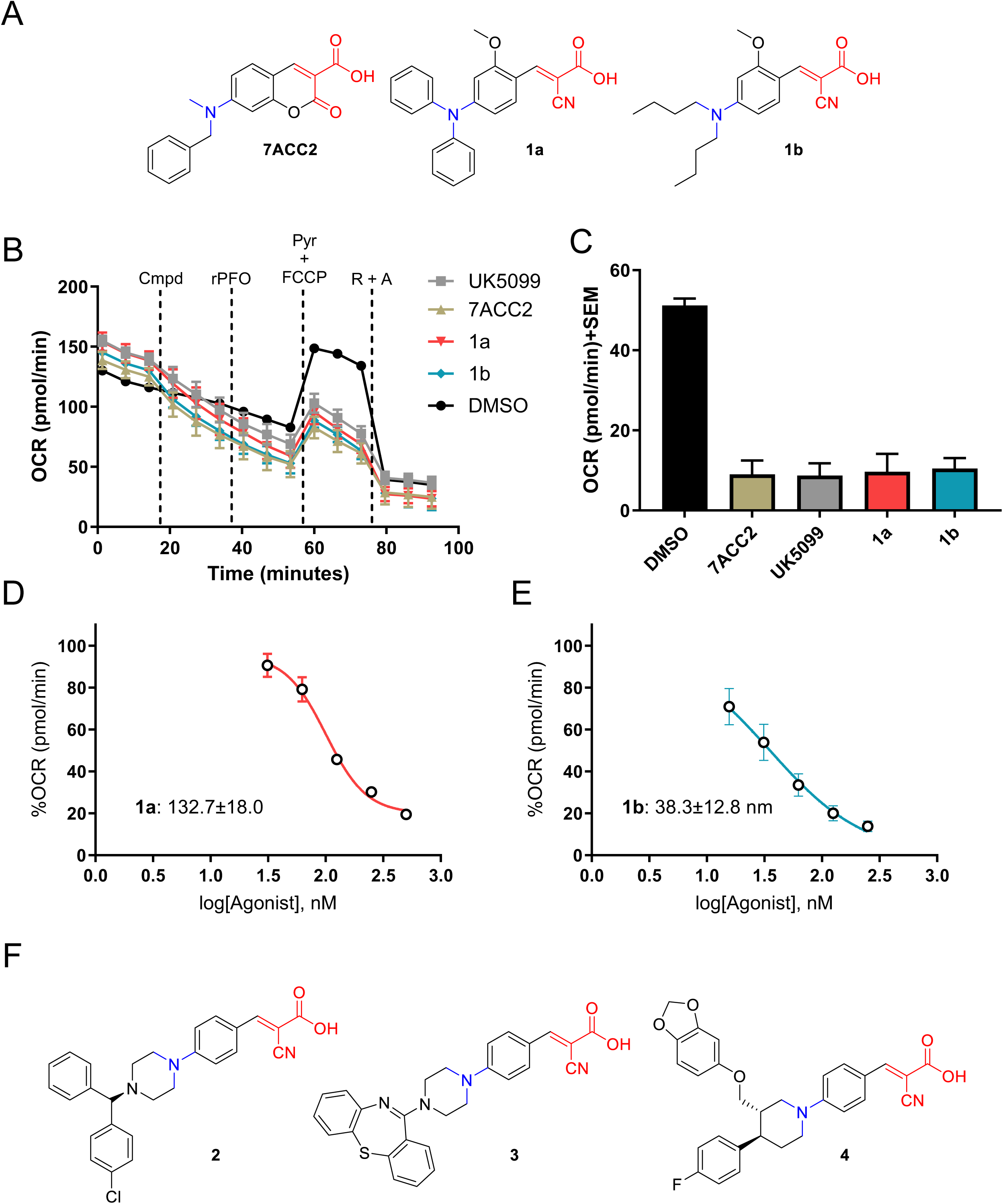
*N,N*-dialkylcinnamic acid-based compounds inhibit the MPC. (**A**) Structure of known MPC inhibitor 7ACC2, and our previously reported MCT1/4 inhibitors **1a** and **1b**. Note structural similarities highlighted in red and blue. (**B**) MPC inhibition assay of candidate compounds **1a** and **1b** in permeabilized 4T1 cells. Acute changes in oxygen consumption rates following the successive injections of respective inhibitors (1 µM), rPFO (1 nM), oligomycin (oligo, 1µM), pyruvate (5mM) + FCCP (0.125µM), and rotenone + antimycin a (Rot/AA, 0.5µM) allow for the calculation of mitochondrial respiratory parameters maximal respiration including pyruvate driven maximal respiration, as highlighted in (**C**). Dose titration of (**D**) **1a** and (**E**) **1b** as calculated from pyruvate driven maximal respiration experiments highlighted in panel (**B**). (**F**) Structures of synthesized *N,N*-dialkylcinnamic acid-based compounds **2-4** on pharmaceutically privileged templates.

### Synthesis of N-piperazinyl and N-piperidinyl cyanocinnamic acids **2-4**

The synthesis of *N*-piperazinylcyanocinnamic acid-based drug candidates **2-4** was carried out in three similar but independent schemes based on varying functionalized piperazine/piperidine substrates (**Scheme 1A-C**). Synthesis of **2** was initiated using standard nucleophilic aromatic substitution of (*S*)-1-((4-chlorophenyl)(phenyl)methyl)piperazine **2a** onto 4-fluorobenzaldehyde under basic conditions to obtain (*S*)-4-(4-((4-chlorophenyl)(phenyl)methyl)piperazin-1-yl)benzaldehyde **2b**, which was subjected to Knoevenagel condensation to obtain (*S*)-*E*-3-(4-(4-((4-chlorophenyl)(phenyl)methyl)piperazin-1-yl)phenyl)-2-cyanoacrylic acid **2** (**Scheme 1A**). Similar reaction schemes were utilized for **3** and **4** wherein 11-(piperazin-1-yl)dibenzo[b,f][1,4]thiazepine **3a** and (3*S*,4*R*)-3-((benzo[d][1,3]dioxol-5-yloxy)methyl)-4-(4-fluorophenyl)piperidine **4a**, respectively, underwent nucleophilic aromatic substitution onto 4-fluorobenzaldehyde to obtain the corresponding aldehydes **3b** and **4b**, respectively. Knoevenagel condensation was performed on the intermediate aldehydes **3b** and **4b** to obtain the corresponding products (*E*)-2-cyano-3-(4-(4-(dibenzo[b,f][1,4]thiazepin-11-yl)piperazin-1-yl)phenyl)acrylic acid **3** and (*E*)-3-(4-((3*S*,4*R*)-3-((benzo[d][1,3]dioxol-5-yloxy)methyl)-4-(4-fluorophenyl)piperidin-1- yl)phenyl)-2-cyanoacrylic acid **4**, respectively.

### MTT cell proliferation inhibition of cyanocinnamic acid derivatives **2-4**

We evaluated compounds **2-4** for their cell proliferation inhibition properties using MTT assay in MDA-MB-231, WiDr, 4T1, MIAPaCa-2, MCF7, and 67NR cell lines. These studies revealed that compounds exhibited good potency across the cell lines tested with IC_50_ values ranging from 10-85 µM for **2**, 7-91 µM for **3**, and 5-73 µM for **4** (**Figure 2A-G**). Interestingly, these compounds exhibited heightened cell proliferation inhibition in 4T1 and WiDr cell lines that have been shown to exhibit high basal levels of mitochondrial respiration (**Figure 2C&D**). These results indicated that anticancer properties of candidate compounds may depend on the cancer cells basal metabolic phenotype, directing their utility toward mitochondrial-active cell lines and consistent with the hypothesized mechanism of action as MPC inhibitors, and consistent with observations reported on our aminocarboxycoumarin candidates^22^.

**Figure 2.**
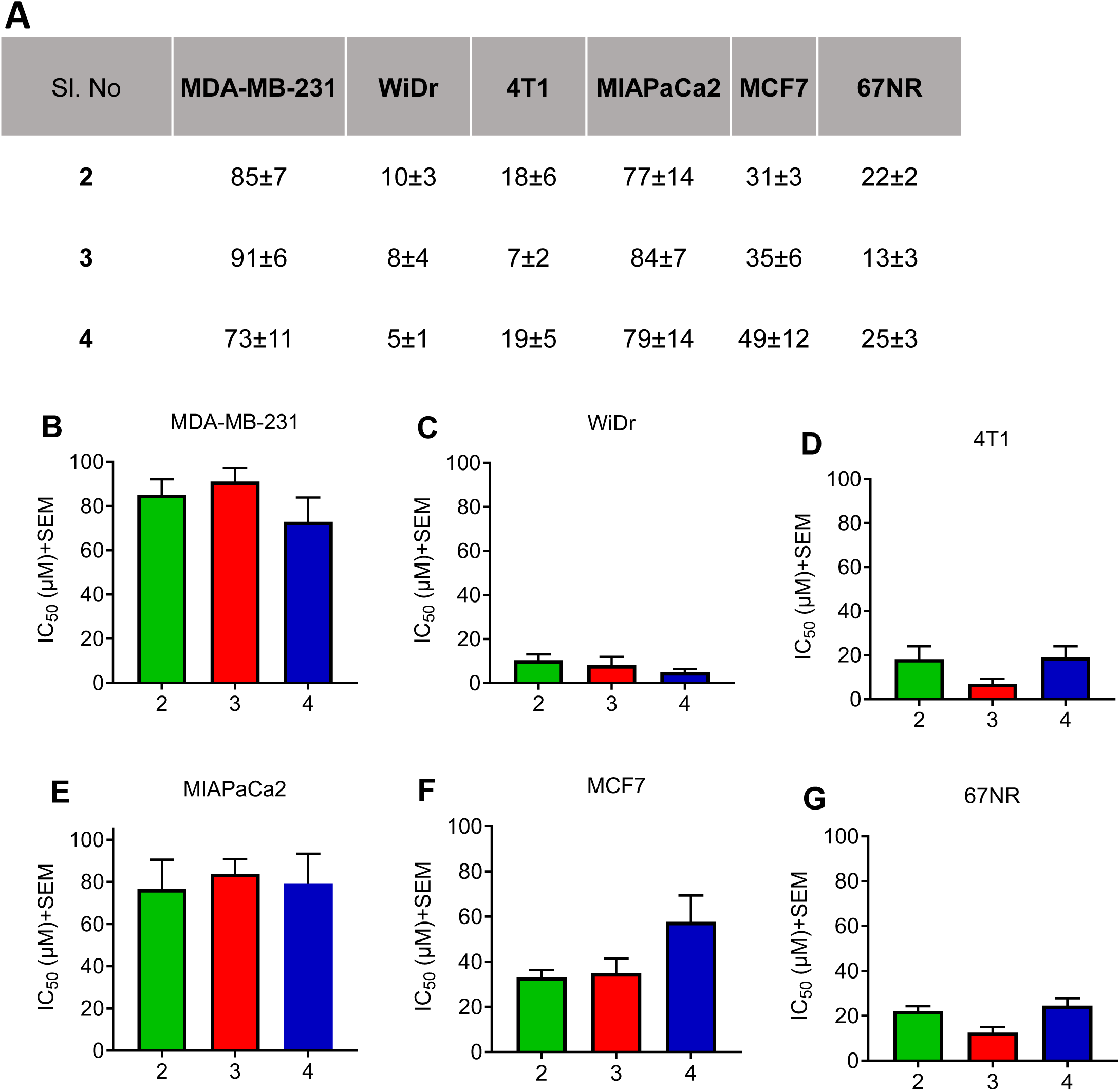
(**A**) IC_50_ values (*µM) of *in vitro* cell proliferation inhibition properties of compounds **2-4** in (**B**) MDA-MB-231, (**C**) WiDr, (**D**) 4T1, (**E**) MIAPaCa-2, (**F**) MCF7, and (**G**) 67NR cells. *IC_50_ values represent the average ± SEM of at least three independent experiments (n=3 biological replicates), with duplicate concentration measurements per experiment (n=2 technical replicates).

### Mitochondrial stress test of cyanocinnamic acid derivatives **2-4**

Treatment with an MPC inhibitor may result in different effects depending on a combination of basal metabolic phenotype and metabolite transporter expression levels. In this regard, Seahorse XFe96 based mitochondrial stress test for cyanocinnamic acids **2-4** was performed in MDA-MB-231, WiDr, and 4T1 cells. These studies illustrated that treatment with compounds **2-4** induced acute respiratory dysfunction in all of the cell lines tested (**Figure 3A-F**). Specifically, in the presence of inhibitor alone, an acute decrease in OCR was observed, indicating a direct effect of compound treatment on mitochondrial respiration (**Figure 3A-C**). Further, following oligomycin treatment, compounds **2-4** inhibited ATP synthesis most substantially in mitochondrial-fueled cell lines WiDr and 4T1 (**Figure 3E-F**). Compounds **2-4** completely suppressed the FCCP-stimulated maximal respiration, potentially indicating the inability of these cells to oxidize mitochondrial fuel consistent with MPC inhibition (**Figure 3D-F**). Treatment with **2-4** induced proton leak in the cell lines tested, also consistent with the inability of a proton-motive force to drive ATP synthesis as mentioned previously (**Figure 3D-F**). A complete loss of the cells ability to increase metabolic rates in response to FCCP-stimulated metabolic dysfunction was observed, indicative of dysfunctional pyruvate flux and MPC inhibition. In this regard, the acute respiratory effects of **2-4** treatment did not depend on basal mitochondrial respiration and general mitochondrial dysfunction was observed in MDA-MB-231, WiDr, and 4T1 cell lines.

**Figure 3.**
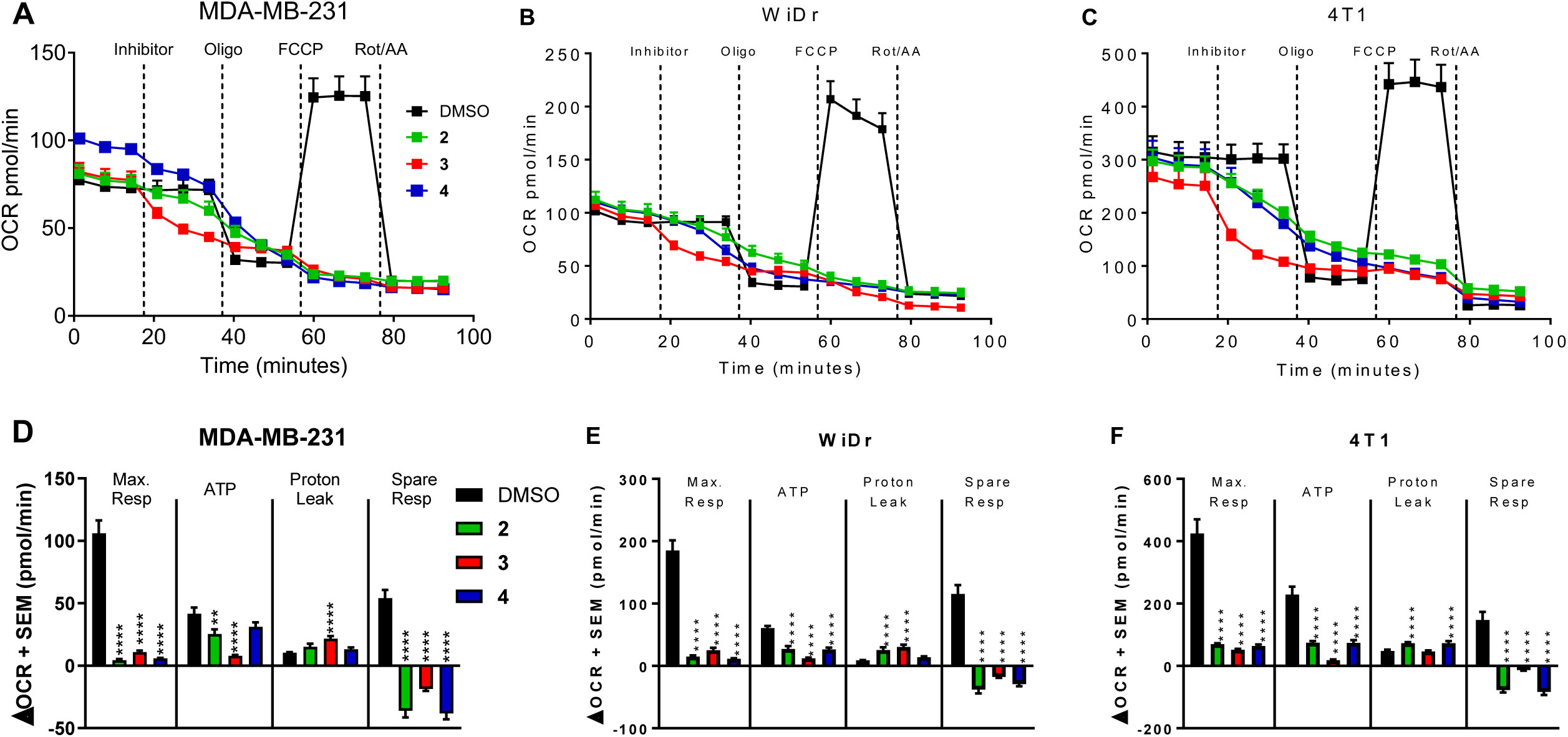
Seahorse XFe96 based mitochondrial stress test in (**A**) MDA-MB-231, (**B**) WiDr, and (**C**) 4T1 cells. (**D-F**) Acute changes in oxygen consumption rates following the successive injections of inhibitor (**2-4**, 30µM), oligomycin (oligo, 1µM), FCCP (MDA-MB-231, 0.25µM; WiDr, 1µM; 4T1, 0.125µM), and rotenone + antimycin a (Rot/AA, 0.5µM) allow for the calculation of mitochondrial respiratory parameters maximal respiration (max resp.), ATP production (ATP), proton leak, and spare respiratory capacity (spare resp.) Bar graphs represent the average ± SEM of at least three separate experiments with n=6 technical replicates in each experiment. One-way ANOVA analysis was performed to evaluate statistical significance between compound **2-4** treated cells versus the control (\*\**p*<0.01, \*\*\*\**p*<0.0001).

### MPC inhibition of cyanocinnamic acid **3** in permeabilized 4T1 cells

Due to the enhanced 4T1 cell proliferation inhibition properties (**Figure 2D**) and substantial acute respiratory response observed in mitochondrial stress tests on this cell line (**Figure 3C**), compound **3** was selected as a lead derivative. Seahorse XFe96 based methods were again employed with slight modifications in permeabilized 4T1 cell line. Recombinant perfringolsyin-O (rPFO) is a cholesterol-dependent secretory cytolysin from Clostridium perfringins that specifically forms pores in the plasma membrane of cells (leaving organelle membranes intact) allowing the selective passage of metabolites; bypassing membrane-transporter mediated function^23^. Use of rPFO in this regard allows for acute delivery of specific metabolites to the mitochondria in whole cells, and corresponding OCR can be observed to evaluate the inhibitory capacity of lead compound **3** toward specific metabolites. An acute decrease in OCR was observed in compound **3** treated 4T1 cells when compared to DMSO control cells (**Figure 4A**). Compound **3**-treated cultures led to a substantial decrease in the pyruvate driven maximal respiration, similar to that of known MPC inhibitor UK5099 (**Figure 4A&B**).

**Figure 4.**
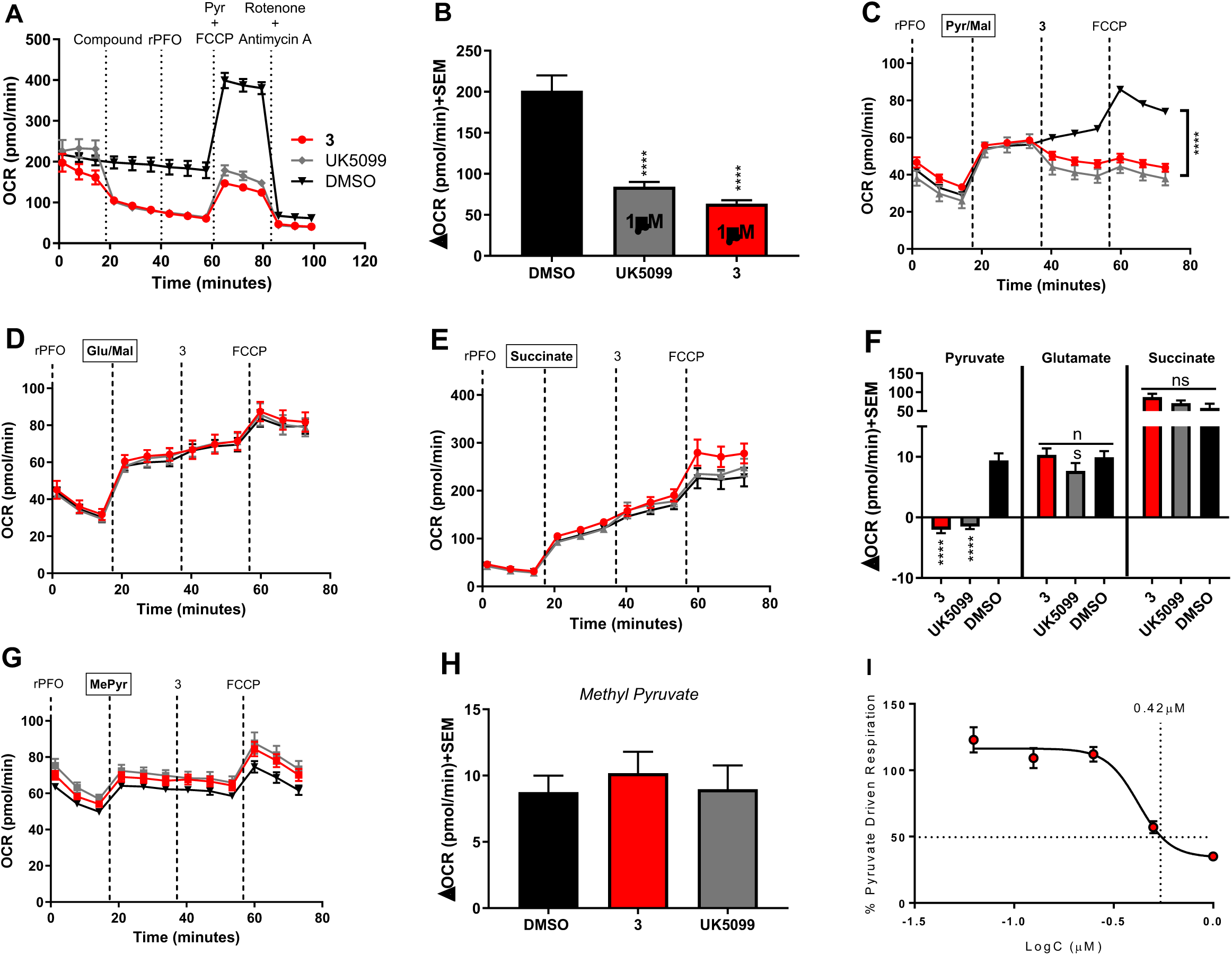
Compound **3** inhibits the mitochondrial pyruvate carrier. (**A-F**) Compound **3** acutely and specifically inhibits pyruvate driven respiration in permeabilized 4T1 cells without affecting glutamate or succinate driven respiratory processes. (**G-H**) Methyl pyruvate reverses the inhibitory capacity of compound **3** toward pyruvate driven respiration, indicating MPC targeting. (**I**) Compound **3** dose response experiments reveal dose-dependent inhibitory capacity of mitochondrial pyruvate uptake.

To further evaluate the mode of action of **3**, permeabilized 4T1 cells were offered metabolic substrates pyruvate (Pyr/Mal **Figure 4C**), glutamate (Glu/Mal, **Figure 4D**), or succinate (**Figure 4E**), followed by the treatment with compound **3** or UK5099 and FCCP and the resulting OCR was observed. The acute changes in OCR demonstrated that compound **3** and UK5099 specifically inhibited pyruvate driven respiration without affecting glutamate or succinate driven respiratory processes (**Figure 4F**). To support that inhibition of pyruvate respiration is due to mitochondrial pyruvate uptake and not due to other pyruvate processing enzymes such as pyruvate dehydrogenase, a parallel experiment using methyl pyruvate (MePyr) was employed (**Figures 4G&H**). MePyr is permeable to the mitochondrial membranes, where in the matrix mitochondrial esterase hydrolysis activity provides pyruvate. Hence, MePyr provides a means to deliver pyruvate to the matrix independent of MPC activity, and reversal of the effects on pyruvate driven respiration in this regard demonstrates specific inhibition of MPC by compound **3**. These experiments illustrated that pyruvate driven respiration effects of compound **3** were completely reversed to the same extent as UK5099, indicating that **3** is an acute and specific MPC inhibitor (**Figure 4G&H**). To evaluate the potency of MPC inhibition, cells were exposed to a range of concentrations of **3** to determine the 50% MPC IC_50_ of maximal pyruvate driven respiration, which was found to be 420 nM (**Figure 4I**).

### Glucose and mitochondrial stress tests of MPC inhibitor **3** in combination with GLUT1 inhibitors

The effectiveness of three GLUT1 inhibitors BAY-876, STF31, and WZB117 (**Figure 5A**) towards glycolysis inhibition were evaluated in glycolytic cell line MDA-MB-231 using Seahorse XFe96 based glycolysis stress test. The results indicated that BAY-876 potently inhibited glycolysis at 1 and 10 µM concentrations (**Figures 5B&E**), whereas STF31 and WZB117 did not affect glycolysis at same concentrations (**Figures 5C-E**). Due to the combination of literature reported metabolic stability and oral bioavailability, along with high selectivity over other GLUT isoforms, BAY-876 was selected as the glycolysis inhibiting counterpart to MPC inhibitor **3**. The combination of compound **3** in the presence of BAY-876 was evaluated using mitochondrial stress test, wherein the OCR and ECAR values were measured simultaneously. This study showed that the combination of BAY-876 and **3** effectively and concurrently inhibited mitochondrial respiration and glycolysis, thereby circumventing the metabolic plasticity exhibited by MDA-MB-231 cells (**Figure 5F-H**).

**Figure 5.**
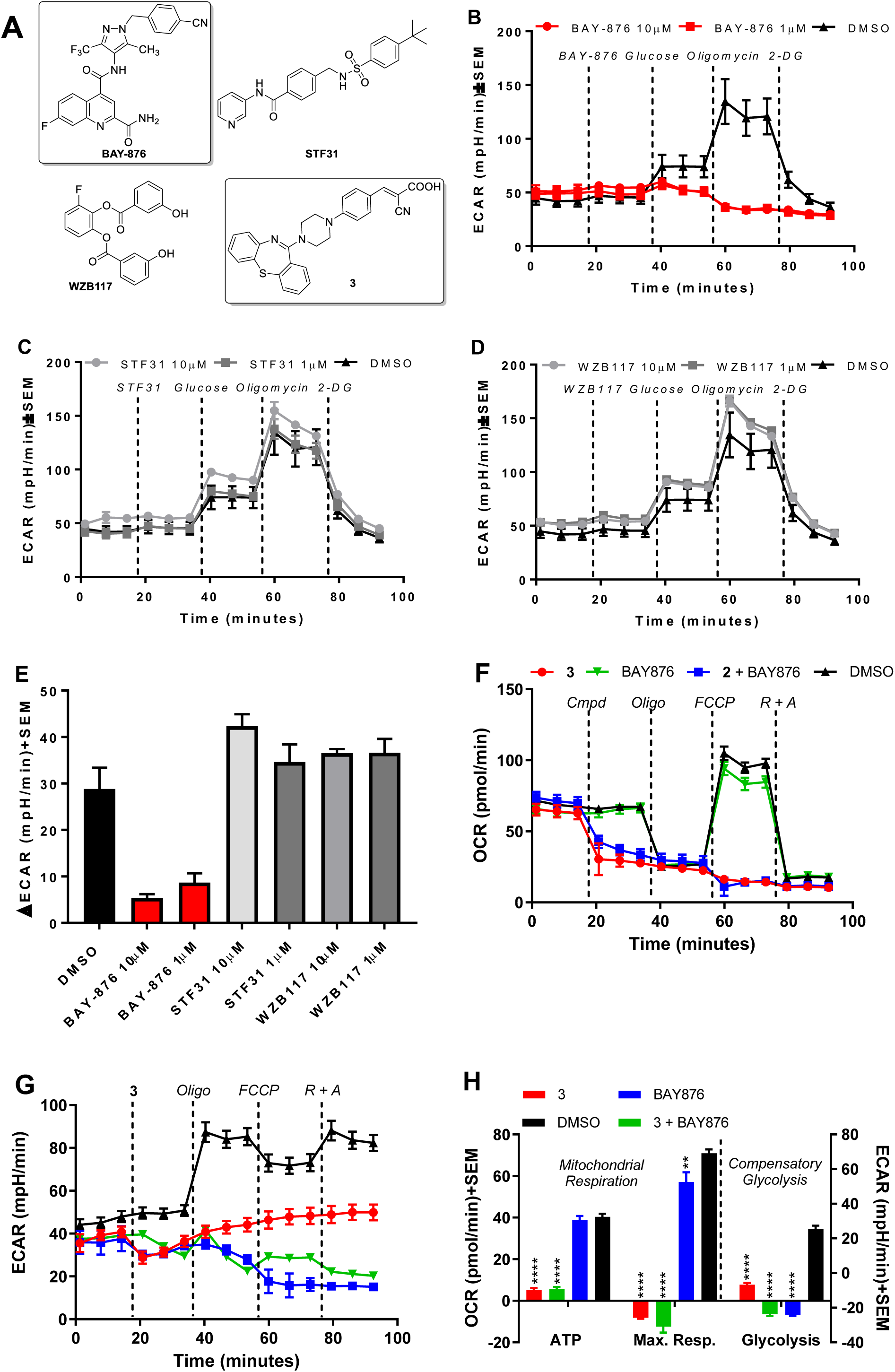
MPC inhibitor **3** in combination with GLUT1 inhibitor BAY-876 diminishes metabolic plasticity in MDA-MB-231 cells. (**A**) Structures of GLUT1 inhibitors BAY-876, STF31, and WZB117, along with MPC inhibitor **3**. (**B, E**) BAY-876 more potently inhibits glycolysis when compared to (**C, E**) STF31 and (**D, E**) WZB117. (**F-H**) MPC inhibitor **3** in combination with BAY-876 inhibits mitochondrial respiration and compensatory glycolysis. Data represents the average± SEM of 5 technical replicates (n=5) of one experiment (N=1). Statistics were evaluated on technical replicates (n=5) and significance was determined using one-way ANOVA, *p<0.05, **p<0.01, ***p<0.001, and ****p<0.0001.

### Synergistic effect of MPC inhibitor **3** in combination with GLUT1 inhibitor BAY-876

Next, the synergistic efficacy of MPC inhibitor **3** in combination with GLUT1 inhibitor BAY-876 was evaluated using cell proliferation inhibition assay in 4T1 cell line, which is known for its aggressive nature and metabolic plasticity. The results showed IC_50_ values of 8, 0.2, and 0.6 µM for compound **3**, BAY-876 and their combination, respectively (**Figure 6A&B**). Isobologram analysis of this combination revealed that this combination strategy is highly synergistic at inhibiting cell proliferation (**Figure 6C**). Thus, MPC inhibitors in combination with GLUT1 inhibitors are deemed a potentially effective combination treatment strategy.

**Figure 6.**
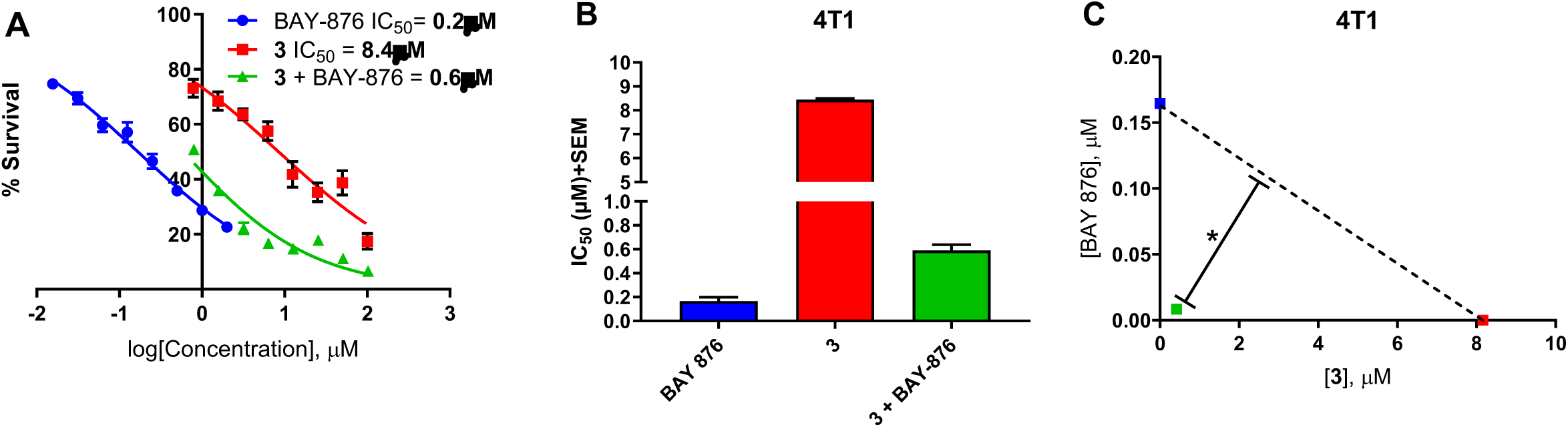
MPC inhibitor **3** and BAY-876 synergize to inhibit breast cancer 4T1 cell proliferation in vitro. (**A, B**) Dose response MTT assay experiments with **3**, BAY-876, and their combination illustrate IC_50_ values of 0.2, 8.4, and 0.6µM respectively. (**C**) Isobologram analysis indicates the combination of **3** and BAY-876 (green) is *synergistic. Dose response curves and IC_50_ values are representative of the average ± SEM of three independent experiments (N=3). Significance based on isobologram was determined as previously described^48^.

## DISCUSSION

The intricate connection between cellular metabolism and proliferation is highly complex and involves crosstalk between both metabolic and cell-cycle machineries.^24-30^ Hence, inhibition of metabolic processes can be reasoned to disrupt cell proliferation. Rapid cell proliferation observed in cancer cells is dependent on both bio-energetic and bio-synthetic thresholds, and control of the cell cycle is highly regulated through energy sensing and biosynthetic mechanisms.^24-30^ Such metabolically relevant sensory mechanisms include but are not limited to 6-phosphofructo-2-kinase/fructose-2,6-biphosphatase 3 (PFKFB3) activity^31^, pyruvate kinase (PKM2) expression^32^, ATP/AMP ratios and AMPK activity^33-35^, NAD+/NADH states through sirtuin deacetylases^36^, and importantly, the availability of acetyl-CoA; a byproduct of numerous metabolic pathways including the tri-carboxylic acid (TCA) cycle^37^. It can be reasoned that potent inhibition of glycolysis and lactate shuttles and disruption of mitochondrial metabolism with compound **3** and BAY-876 may lead to a cellular metabolic crisis that senses cell cycle machinery to stop proliferating during phases of the cell cycle that are dependent on the genetic make-up of the cell. Further, it can be reasoned that the activation of cell death pathways will be dependent on the integrity of the molecular players in these pathways such as important master transcription factor p53. MDA-MB-231 and WiDr are p53 mutant cell lines^38,39^, and responses to cellular and metabolic stress may result in unpredictable cell-cycle arrest and apoptotic responses as p53 is an important mediator between metabolic and proliferative states^40^. Specifically, p53 has been shown to modulate MCT1 status through direct interaction with the MCT1 gene promoter along with alterations in the stability of MCT1 mRNA^41^. Due to the integral relationship of extracellular lactate uptake via MCT1 and mitochondrial pyruvate uptake via MPC, it would not be surprising if dysfunction in MPC and/or GLUT1 would lead to an anti-proliferative response, potentially depending on a combination basal metabolic phenotypes, transporter expression, and p53 status.

Specific molecular mechanisms by which the above-mentioned metabolic cell-cycle regulators function may also play a role in the phase at which the cell cycle is disrupted. For example, glycolytic enzyme PFKFB3 promotes progression through M-phase through activation of mitotic kinase CDK1, phosphatase Cdc25, and down-regulation of M-phase inhibitor p27^31^. Additionally, PKM2 which functions as a molecular shuttle from glycolysis to the TCA cycle, also plays regulatory roles in the cell-cycle by phosphorylating the spindle assembly checkpoint protein Bub3 resulting in proper chromosomal segregation during mitosis^32^. Moreover, the availability of acetyl CoA plays important roles in the cells global ability to turn on transcription and cell-cycle machineries. The glycosylation of proteins, namely transcription factors, by O-linked-N-acetylglucosamine (O-GlcNAc) transferases (OGT) plays an important role in cell proliferation-contingent on availability of acetyl CoA^42-45^. Interestingly, OGT activity is at its highest during the G2/M phase of the cell cycle where it modulates the activity of the cell-cycle checkpoint protein cyclin B. Inhibition of OGT activity in this regard has been shown to decrease cyclin B activity and consequential progression through M-phase^25,45^. Also, OGTs play important roles in the stability of the mitotic spindle through the OGT/OGA/Aurora B/PP1 complex^46^. Further, histone acetylation facilitated by histone acetyl transferase (HAT) plays a vital role in chromatin modifications prior to cell cycle entry, and is limited by the availability of cytosolic acetyl CoA^37,47,48^. In this regard, OGT and HAT activities are crucial molecular players in the cells ability to proliferate, and starvation of acetyl CoA through metabolic dysfunction induced by compound **3** and/or BAY-876 may lead to inhibition of both. Thus, disrupting pathways that are vital in the maintenance of acetyl CoA status such as glycolysis and mitochondrial metabolism (TCA cycle) and pyruvate shuttling with the proposed combination of MPC inhibitor **3** and BAY-876 may limit the cells’ ability to modify chromatin and important transcription factors for the synthesis of a variety of important cell-cycle machineries.

## CONCLUSIONS

In the current study, novel cyanocinnamic acid-based MCT/MPC inhibitors **2-4** were synthesized based on pharmacologically privileged *N*-piperazinyl and *N-*piperidinyl templates. *In vitro* cell proliferation inhibition illustrated that compounds **2-4** inhibited cancer cell proliferation in the low micromolar range, improved when compared to first generation candidates. Seahorse XFe96 based mitochondrial stress tests provided **2-4** potently and acutely inhibited numerous parameters of mitochondrial respiration in MDA-MB-231, WiDr, and 4T1 cells. Additionally, lead candidate compound **3** inhibited pyruvate driven respiration without affecting glutamate or succinate fueled respiratory processes in permeabilized 4T1 cells. To expand on the clinical utility of this inhibitor, combination studies with GLUT1 inhibitor BAY-876 illustrate the capacity of these compounds to inhibit metabolic plasticity in triple negative breast cancer MDA-MB-231 cells and is synergistic in inhibiting cell proliferation in aggressive stage IV breast cancer 4T1. The studies herein provide novel and potent MPC inhibitors that can be potentially utilized as metabolism targeting agents for anticancer therapy.

## MATERIALS AND METHODS

### General chemistry procedures

Commercial grade solvents and reagents were purchased from Fisher Scientific (Houston, TX), Sigma-Aldrich (Milwaukee, WI), or Ambeed and were used without further purification. ^1^H and ^13^C NMR spectra were recorded in the indicated solvent on a Varian Oxford-500 MHz spectrometer at 500 and 125 MHz for ^1^H and ^13^C, respectively. Multiplicities are indicated by s (single), d (doublet), t (triplet), m (multiplet), dd (doublet of doublet) and br (broad). Chemical shifts (δ) are reported in parts per million (ppm) and coupling constants (*J*), in hertz. Elemental analysis (CHN) results were obtained from Atlantic Microlab services.

### Representative synthesis of (4-(4-((4-chlorophenyl)(phenyl)methyl)piperazin-1-yl)phenyl)-2-cyanoacrylic acid **2**

To a stirred solution of (*S*)-1-((4-chlorophenyl)(phenyl)methyl)piperazine (10 mmol) in DMF (20 mL), was added 4-fluorobenzaldehyde (10 mmol) and potassium carbonate (20 mmol) and refluxed for 12 hours. Upon completion of the reaction (TLC), the reaction mixture was extracted 3X with water and ether (50 mL). The organic layer was separated, dried, and evaporated under vacuum to obtain 4-(4-benzhydrylpiperazin-1-yl)benzaldehyde. To a solution of this aldehyde (5 mmol) in acetonitrile (10 mL), piperidine (7.5 mmol) and cyanoacetic acid (7.5 mmol) were added and the reaction mixture was refluxed at 120 °C for 8 hours. Upon the completion of the reaction, the contents were transferred into 6M ice cold HCl and the resulting solid was filtered and washed with water. The crude product was recrystallized in ethylacetate:methanol (5:1) to obtain pure product. Compounds **3** and **4** were synthesized using similar scheme.

### (S,E)-3-(4-(4-((4-chlorophenyl)(phenyl)methyl)piperazin-1-yl)phenyl)-2-cyanoacrylic acid **2**

^1^H NMR (500 MHz, DMSO-d_6_): δ 12.72 (s, 1H), 8.13 (s, 1H), 7.98-7.92 (m, 5H), 7.52-7.36 (m, 5H), 7.06 (d, *J* = 9.0 Hz, 2H), 5.72 (s, 1H), 4.09 (s, 2H), 3.72 (s, 2H), 3.16-3.03 (m, 4H); ^13^C NMR (125 MHz, DMSO-d_6_): δ 172.49, 164.72, 154.08, 152.83, 133.71, 130.93, 129.87, 128.90, 121.73, 117.74, 114.55, 110.00, 109.98, 97.38, 73.68, 50.84, 43.36, 31.14; Anal Calc’d for C_16_H_15_NO_4_ (285.30): C 67.36, H 5.30, N 4.91, found: C 67.38, H 5.19, N 4.98

### (E)-2-cyano-3-(4-(4-(dibenzo[b,f][1,4]thiazepin-11-yl)piperazin-1-yl)phenyl)acrylic acid **3**

^1^H NMR (500 MHz, DMSO-d_6_): δ ^13^C NMR (125 MHz, DMSO-d6): δ 188.8, 172.7, 172.5, 164.9, 161.8, 154.2, 153.2, 140.1, 133.8, 133.4, 133.1, 131.6, 130.8, 130.4, 130.0, 126.6, 120.9, 118.0, 113.7, 110.0, 96.2, 48.2, 45.7; Anal Calc’d for C_16_H_15_NO_4_ (285.30): C 67.36, H 5.30, N 4.91, found: C 67.38, H 5.19, N 4.98

### (E)-3-(4-((3S,4R)-3-((benzo[d][1,3]dioxol-5-yloxy)methyl)-4-(4-fluorophenyl) piperidin-1-yl)phenyl)-2-cyanoacrylic acid **4**

^1^H NMR (500 MHz, DMSO-d_6_): δ 8.09 (s, 1H), 7.95 (d, *J* = 9.5 Hz, 2H), 7.27-7.24 (m, 2H), 7.11-7.06 (m, 4H), 6.72 (d, *J* = 9.0 Hz, 1H), 6.52 (d, *J* = 2.5 Hz, 1H), 6.21 (dd, *J* = 8.8Hz, 2.8 Hz, 1H), 5.91 (s, 2H), 4.32 (d, *J* = 11.5 Hz, 1H), 4.19 (d, *J* = 13.0 Hz, 1H), 3.65 (dd, *J* = 10.0Hz, 1H), 3.55-3.52 (m, 1H), 3.38-3.34 (m, 1H), 3.09-3.01 (m, 2H), 2.89-2.84 (m, 1H), 2.49-2.48 (m, 1H), 2.15-2.11 (m, 2H); ^13^C NMR (125 MHz, DMSO-d_6_): δ 165.11, 162.28, 160.35, 154.36, 154.02, 153.73, 148.30, 141.69, 140.02, 134.12, 129.69, 129.63, 120.05, 118.16, 115.72, 115.56, 113.85, 108.37, 106.09, 101.43, 98.42, 95.23, 69.12, 65.37, 50.23, 47.65, 43.74, 41.13, 33.38; Anal Calc’d for C_16_H_15_NO_4_ (285.30): C 67.36, H 5.30, N 4.91, found: C 67.38, H 5.19, N 4.98

### Cell lines and culture conditions

MDA-MB-231 cells (ATCC) were grown in DMEM medium supplemented with FBS (10%) and penicillin-streptomycin (50U/ml, 50µg/ml). MCF7 cells (ATCC) were grown in α-MEM with FBS (5%), non-essential amino acids (0.1 mM), insulin (10 µg/mL), sodium pyruvate (1 mM), epidermal growth factor (100 ng/mL), hydrocortisone (10 µg/mL), HEPES (10 mM), and penicillin-streptomycin (50U/ml, 50µg/ml). MIAPaCa-2 cells (ATCC) were grown in DMEM medium supplemented with FBS (10%), horse serum (2.5%) and penicillin-streptomycin (50U/ml, 50µg/ml). WiDr cells (ATCC) were cultured in MEM media supplemented with FBS (10%) and penicillin-streptomycin (50U/ml, 50µg/ml). 4T1 and 67NR cells (ATCC) were grown in RPMI medium supplemented with FBS (10%) and penicillin-streptomycin (50U/ml, 50µg/ml).

### Cell proliferation inhibition assay

Cell proliferation inhibition properties were evaluated using MTT assay. Briefly, cells were seeded in 96 well plates (5×10^3^ cells/well) where they were incubated to adhere for 12-24 hours. Compounds were then added to the wells in serial dilution for a total of eight concentrations starting from 100 µM concentration. DMSO was used to dissolve stock solutions at 1000X of starting concentration, resulting in DMSO cell concentration of 0.1% (v/v). Cells were then incubated in test compound for 72 hours. MTT (0.5 mg/ml in 1X PBS) was added and incubated for four hours at 37°C. Ensuing formazan precipitate was dissolved using sodium dodecyl sulfate solution (0.1g/L in 0.01N HCl) and incubated for further four hours. Absorbance at 570 nm was recorded and IC_50_ values of compounds were calculated using the absorbance of untreated wells as 100% survival via GraphPad Prism software. Synergy of combination studies was evaluated using compusyn software as previously described^48^.

### Mitochondrial and glycolysis stress tests

These assays were conducted in accordance with previously reported methods^20^.

### MPC inhibition in permeabilized 4T1 cells

Permeabilized cell assays were performed using rPFO as previously described^22-23^ with slight modifications. 4T1 cells were seeded (20,000cells/well) onto Seahorse XFe96 well plates and incubated overnight in growth media at 37 °C and 5% CO_2_ for adherence. On the day of the assay, growth media was aspirated and replaced with mannitol/sucrose buffer (MAS; 70mM sucrose, 220mM mannitol, 10mM potassium phosphate monobasic, 5mM magnesium chloride, 2mM HEPES, and 1mM EGTA) after 3X rinse of growth media to remove serum and endogenous metabolic substrates and incubated at 37 °C in a non-CO_2_ incubator. Respective inhibitor and substrate milieu’s were prepared in MAS buffer for port injections A-D at 8X, 9X, 10X, and 11X the target cell concentrations to account for intrinsic dilution factor of *in situ* injections of each port. For some experiments (**Figure 4A&I**), test compound (1µM) was injected in port A, followed by rPFO (1nM) in port B, followed by respective substrate cocktails (FCCP stimulated) in port C, and rotenone and antimycin A (0.5µM) in port D. In other experiments (**Figure 4C-H**), permeabilization was initiated prior to port A injection during the MAS buffer wash phase, followed by substrates, test compound (1µM), and FCCP (0.125 µM). Final substrate concentrations for specific tests were as follows: (5mM pyruvate, 0.5mM malate, 2mM dichloroacetate (DCA); 10mM glutamate, 0.5mM malate, 2mM DCA; 10mM succinate, 2µM rotenone; 20mM methyl pyruvate, 5mM pyruvate, 0.5mM malate, and 2mM DCA).

## SCHEME LEGENDS

**Scheme 1.**
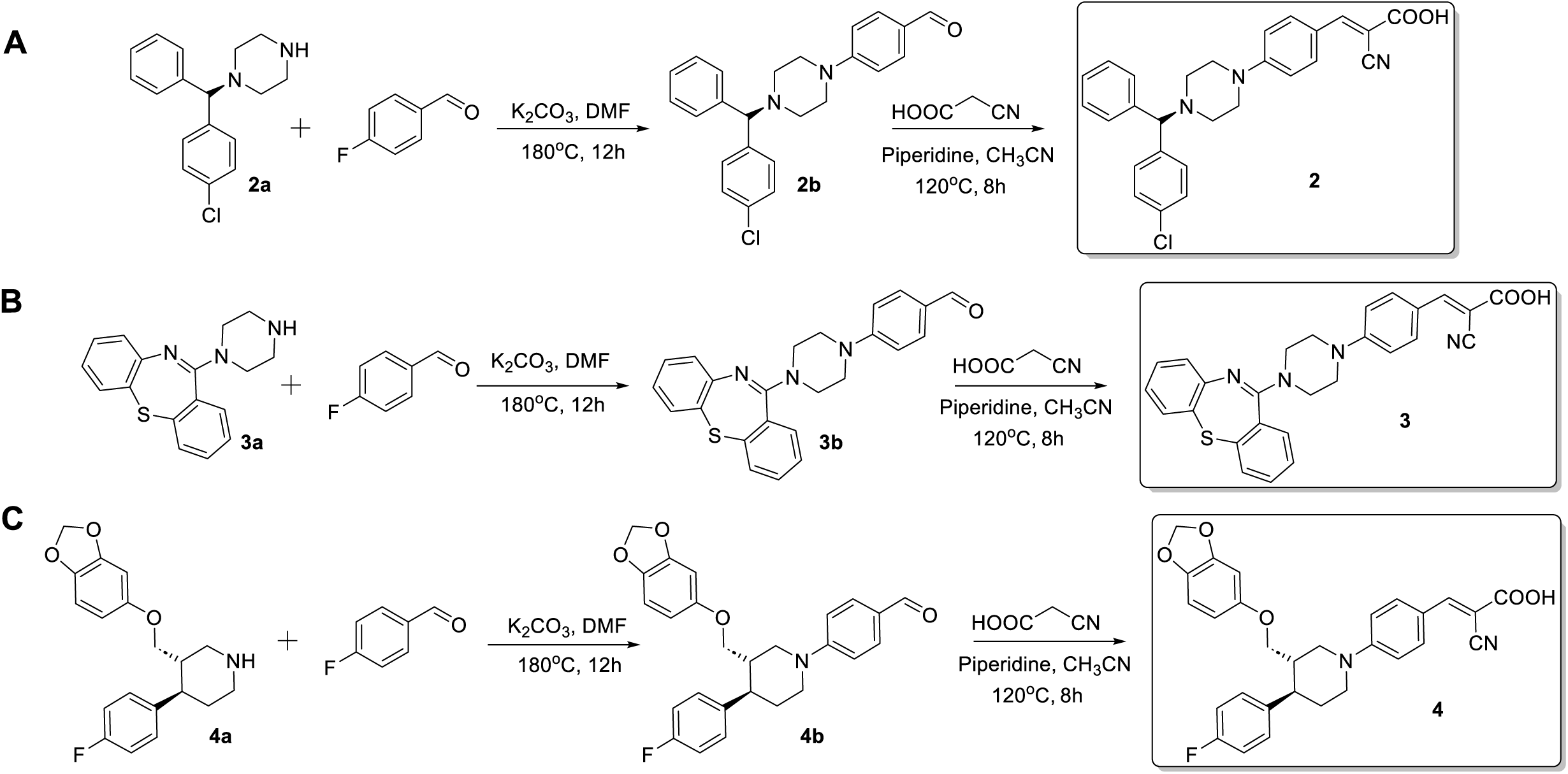
Synthesis of functionalized *N*-piperazinyl and *N*-piperidinyl cyanocinnamic acids **2-4**.

